# MED26-enriched condensates drive erythropoiesis through modulating transcription pausing

**DOI:** 10.1101/2024.08.26.609654

**Authors:** Shicong Zhu, Xiaoting Zhang, Na Li, Xinying Zhao, Man Li, Si Xie, Qiuyu Yue, Yunfeng Li, Dong Li, Fan Wu, Zile Zhang, Ziqi Feng, Yiyang Zhang, Wonhyung Choi, Xinyi Jia, Yuelin Deng, Qi Hu, Xingyun Yao, Xiaofei Gao, Hsiang-Ying Lee

**Affiliations:** Ministry of Education Key Laboratory of Cell Proliferation and Differentiation, School of Life Sciences, Peking University, Beijing 100871, China; Peking-Tsinghua Center for Life Sciences, Academy for Advanced Interdisciplinary Studies, Peking University, Beijing, 100871, China; Key Laboratory of Growth Regulation and Translational Research of Zhejiang Province, School of Life Sciences, Westlake University, Hangzhou, 310030, Zhejiang, China

## Abstract

The Mediator complex regulates various aspects of hematopoietic development, but whether composition of the Mediator complex undergoes dynamic changes for diversifying transcription and functional outputs is unknown. Here, we found that MED26, a subunit in the core Mediator complex, played a distinctive role in facilitating transcription pausing essential for erythroid development. While most Mediator subunits drastically decreased during this process, MED26 remained relatively abundant. Intriguingly, in the early stages, more than half of MED26 occupancy sites did not co-localize with MED1, a representative Mediator subunit, suggesting these subunits exert context-dependent gene regulation. We revealed that MED26-enriched loci were associated with RNA polymerase Ⅱ pausing. MED26 manifested a markedly preferential recruitment of pausing-related factors, leading to an increase in Pol Ⅱ pausing critical for genome-wide transcription repression during erythropoiesis. Moreover, MED26 exhibited pronounced condensate-forming capability, which was necessary for its function in promoting erythropoiesis and recruiting pausing-related factors. Collectively, this study provides mechanistic insights into the functional coordination of distinct Mediator subunits during development and highlights the switch of transcription condensates towards a MED26 enriched form, which modulates transcription pausing to facilitate transcription repression and erythroid development.

## Introduction

During mammalian erythroid development, erythroblasts undergo chromatin reorganization that involves drastic nuclear condensation and global transcription repression before enucleation^1–3^. Terminal erythropoiesis has been associated with declines in histone marks involved in transcription elongation, including H3K36me2, H3K36me3, and H3K79me2^4,5^, along with an increase in H4K20me, a modification associated with Pol Ⅱ pausing and erythroblast chromatin condensation^6,7^. The positive transcription elongation factor (P-TEFb), which contains the catalytic subunit cyclin-dependent kinase 9 (CDK9), cooperates with the master erythroid transcription factor GATA1 to enhance transcription elongation^8^. HEXIM1, a regulator that promotes Pol Ⅱ pausing, is highly expressed during terminal erythropoiesis and is associated with accelerated erythroid differentiation^9^. Collectively, these findings suggest transcription pausing and elongation play regulatory roles in erythropoiesis, yet the mechanisms governing their upstream regulation remain largely unknown.

Transcription pausing is a prevalent regulatory mechanism when RNA polymerase Ⅱ (Pol Ⅱ) initiates transcription but stalls at 20∼60 bp downstream of the transcription start site^10,11^. Pol Ⅱ pausing exerts a pivotal influence on cell fate determination and development by intricately regulating gene expression dynamics, with paused Pol Ⅱ near critical genes acting as molecular sensors, awaiting external cues to swiftly trigger or inhibit transcription and thereby orchestrating precise gene expression patterns pivotal for differentiation^12–14^. The importance of transcription pausing has been demonstrated in drosophila mid-blastula transition and cell specification in mouse embryonic stem cells^15^. Nonetheless, many mechanistic questions regarding Pol Ⅱ pausing in mammalian development remain unanswered, particularly beyond cell specification. For instance, how does pausing impact upon the subsequent differentiation progression?

The Mediator complex is a large multi-subunit complex composed of head, middle, tail, and CDK8 kinase modules and is conserved from yeasts to metazoans^16,17^. The Mediator complex forms a functional bridge between gene promoters and enhancers, linking tissue-specific transcription factors (TFs) with general transcription factors (GTFs) and Pol Ⅱ, thereby serving as an integrative hub for pre-initiation complex assembly, transcription elongation, and termination^16,18^. Several Mediator subunits was shown to be important for erythropoiesis, including MED1 and MED25^19–21^. Previous studies on Mediator-regulated developmental processes have often focused on the function of a single subunit and its cooperation with a TF; however, it remains unclear how individual Mediator subunits functionally cooperate across developmental stages, thereby contributing to the establishment of a context-dependent transcriptional program^22^.

Here, we used erythropoiesis as a paradigm to address the mechasims of global transcription repression and how individual Mediator subunits cooperatively diversify the function of the Mediator complex in a developmental scenario. Our study reveals that while MED1 and MED26 co-occupy a plethora of genes important for cell survival and proliferation at the HSPC stage, MED26 preferentially marks erythroid genes and recruits pausing-related factors for cell fate specification. Gradually, MED26 becomes the dominant factor in shaping the composition of transcription condensates and transforms the chromatin towards a repressive yet permissive state, achieving global transcription repression in erythropoiesis. This study not only unveils the mechanism of Mediator regulating transcription pausing but also how the increasing dominance of a single subunit drives the transformation of transcription states and progression of a developmental process.

## Results

### The levels of most Mediator subunits decreased, while MED26 remained relatively abundant, during erythropoiesis

During terminal erythropoiesis, global transcription repression occurs concurrently with erythroid differentiation, but it remains unclear how global genes are repressed, while few selective genes remain actively transcribed. As the Mediator complex is crucial in the regulation of various transcription steps, we detected its expression in primary human erythroblasts derived from CD34^+^ hematopoietic stem and progenitor cells^23^. We found that the protein levels of most Mediator subunits decreased substantially in the terminal erythroid differentiation; in contrast, unexpectedly, that of MED26 remained detectable throughout (**Figure 1A-1C**). Consistently, immunofluorescence imaging of primary erythroblasts isolated from E14.5 mouse fetal livers revealed that the MED1 level declines while MED26 remains detectable from early to late erythropoiesis (**Figure 1D**). Given the relative abundance of MED26 in terminal erythropoiesis and the critical function of the Mediator complex in transcription regulation, we reason that MED26 may serve a unique function in late erythropoiesis^24^.

**Figure 1.**
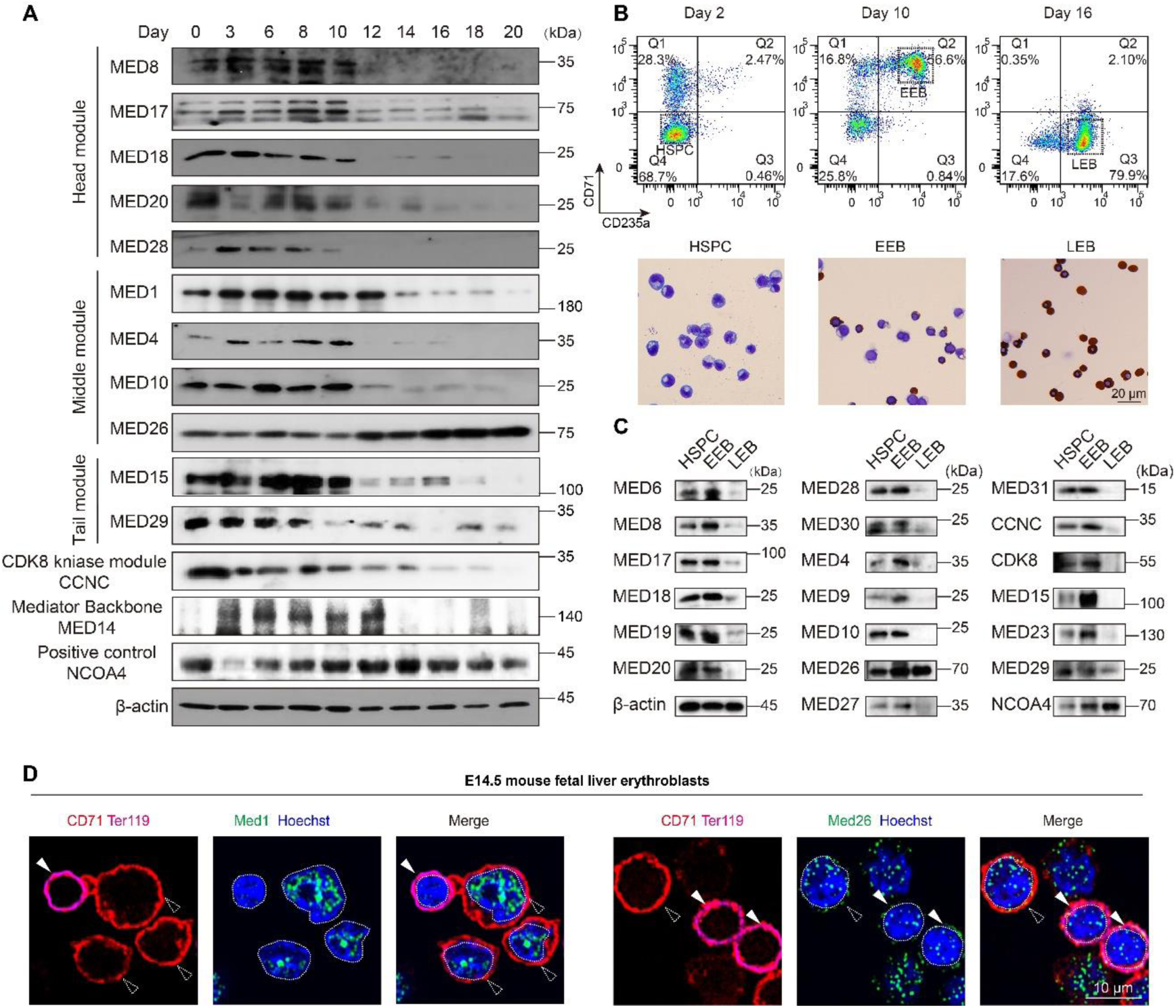
Differential stoichiometry changes of Mediator complex subunits during erythropoiesis. **(A)** Western blot showing the relative abundance of Mediator subunits over time (Days 0-20) in primary human CD34^+^ erythroid culture. β-actin was used as a loading control, and NCOA4, which is highly expressed in terminal erythropoiesis, served as a positive control. **(B)** FACS sorting strategy for isolating indicated cell populations and corresponding micrographs by benzidine-Giemsa staining. **(C)** Western blot showing the relative abundance of Mediator subunits in FACS-sorted cell populations representing different developmental stages (HSPC: CD71^-^CD235a^-^; early erythroblasts, EEB: CD71^+^CD235a^+^; late erythroblasts, LEB: CD71^-^CD235a^+^) in human CD34^+^ erythroid culture with β-actin as a loading control, and NCOA4 as a positive control. **(D)** Immunofluorescence imaging of MED1 (left) or MED26 (right) in mouse fetal liver erythroblasts. CD71^+^Ter119^-^ cells are early erythroblasts (shown by dotted arrows), and CD71^+^Ter119^+^ cells are late erythroblasts (shown by white solid arrows). Hoechst 33342 staining was used to indicate the nuclear area.

### MED26 plays a crucial role in erythropoiesis

To investigate the regulatory role of *Med26* in erythropoiesis *in vivo*, we constructed *Med26* conditional knockout mice with Cre recombinase expression driven by the erythropoietin receptor (EpoR) promoter (*Med26*^flox/flox^; EpoR-Cre/+). The results show that *Med26*-erythroid-specific knockout mice exhibit pale embryos and severe anemia at E14.5 (**Figure 2A**), demonstrating the direct regulatory role of *Med26* in fetal erythropoiesis.

**Figure 2.**
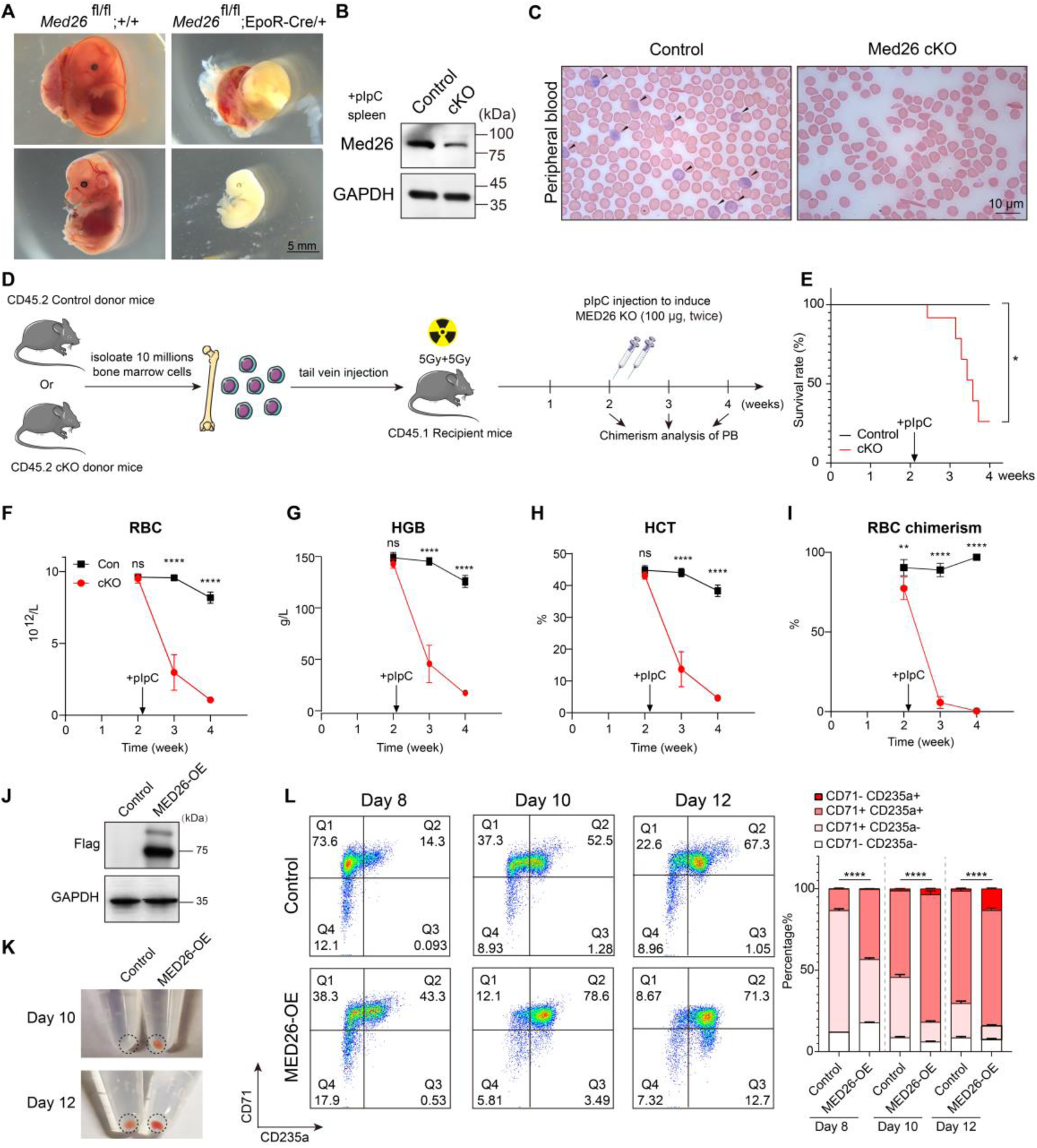
MED26 regulates all stages of erythropoiesis. **(A)** Representative images of mouse embryos of *Med26*^flox/flox^;wt/wt and *Med26*^flox/flox^;EpoR-Cre/wt at E14.5. Scale bar=5 mm. **(B-I)** Analyses conducted in *Med26*^flox/flox^;Mx1-Cre/wt mice. In these panels, “cKO” refers to *Med26*^flox/flox^;Mx1-Cre/wt mice. **(B)** Western blot demonstrating *Med26* knockout efficiency in mouse spleen cells with GAPDH as a loading control. **(C)** Giemsa staining of peripheral blood smears from control and cKO mice. The arrow indicates reticulocytes. For panels (B and C), tests and phenotypes were analyzed 3 days after administering 10 µg/g pIpC**. (D)** Timeline of the mice bone marrow transplantation procedure. Bone marrow cells were collected from CD45.2+ control or cKO mice. Subsequently, 10 million cells were injected into 10Gy (5Gy+5Gy) irradiated myeloablative CD45.1+ recipient mice. After a two-week period of hematopoietic reconstitution, the mice received two injections of 100 µg pIpC (totaling 200 µg) to induce Med26 knockout. Peripheral blood samples were collected weekly from weeks 2-4 for CBC analysis and chimera percentage assessment. **(E)** Survival rate of recipient mice after Med26 knockout. The p value was calculated by Kaplan-Meier log rank test, *, p < 0.05. (C) Giemsa staining of peripheral blood smears from control and cKO mice. Scale bar=10 µm. **(F-I)** The graphs depict the dynamics of erythroid reconstitution before and after Med26 knockout. Data are represented as the mean ± SD (Control group: n=6; cKO group: n=9). Statistical significance is denoted as n.s., not significant, *, p < 0.05, **, p < 0.01, ***, p < 0.001, ****, p < 0.0001 as determined by Student’s t-test. **(J)** Western blot demonstrating MED26 overexpression efficiency in the *ex vivo* human CD34^+^ erythroid differentiation system on Day 8 with GAPDH as a loading control. **(K)** Pictures of CD34^+^ cell pellets upon MED26 overexpression on Day 10 and Day 12. The dotted circles indicate the cell pellet positions. **(L)** FACS analysis to detect two erythroid markers (CD71 and CD235a) upon MED26 overexpression. Data are represented as the mean ± SD (n=3).

Next, to examine the role of *Med26* in adult erythropoiesis, we generated inducible conditional knockout mice (*Med26*^flox/flox^; Mx1-Cre/+). Using this system, pIpC injection induces *Med26* genomic loci recombination at the specified developmental timepoint (**Fig. S1A-B**), leading to substantial downregulation of Med26 protein expression (**Figure 2B**). Giemsa staining of the peripheral blood smear showed the presence of irregularly shaped RBCs and scarce reticulocytes (**Figure 2C**). We further assessed the erythroid recovery capacity differences of *Med26* knockout under stress conditions by irradiation protection and phenylhydrazine induced acute hemolytic anemia models. First, we harvested 10 million bone marrow cells from either control or cKO CD45.2^+^ mice and transplanted them into irradiated myeloablative CD45.1^+^ recipient mice. Following a 2-week period for hematopoietic reconstitution, *Med26* was knocked out via pIpC induction, and CBC analyses were performed weekly (**Figure 2D**). The results showed that prior to *Med26* knockout through pIpC induction, red blood cells in both the control and *Med26* cKO groups were reconstructed to similar levels. However, following the *Med26* knockout, there was a worse survival rate (**Figure 2E**) and a marked decline in erythroid parameters and the chimerism percentage of red blood cells (**Figure 2F-I and Fig. S2**). Additionally, under phenylhydrazine (PHZ)-induced acute hemolytic anemia conditions, cKO mice were unable to recover after PHZ treatment (**Fig. S3**). These results demonstrate that Med26 is essential for erythrocyte reconstitution. Collectively, these findings show that Med26 is crucial for erythropoiesis under both normal and stress conditions.

Furthermore, we conducted a gain-of-function assay in the primary human erythroid culture system^23^, which showed that MED26 overexpression accelerated erythroid differentiation (**Figure 2J-2L**). To determine whether the acceleration is stage-specific, we isolated cells at different stages—HSPCs (CD71^-^CD235a^-^), erythroid progenitors (CD71^+^CD235a^-^) and erythroid precursors (CD71^+^CD235a^+^)—from human CD34^+^ erythroid cultures by FACS sorting and induced MED26 overexpression. Our findings demonstrate that MED26 can accelerate erythropoiesis across all examined stages (**Fig. S4**). In summary, we demonstrate that MED26 is essential for erythropoiesis and that MED26 overexpression can accelerate the process.

### MED26-enriched chromatin sites are associated with RNA polymerase II pausing

To further dissect the molecular basis underlying MED26 function, we analyzed the CUT&Tag signals of MED26 and compared them to those of MED1, which is often regarded as a representative subunit of Mediator, in human CD34^+^ erythroid culture cells on Day 4. Unexpectedly, we found that ∼60% of chromatin sites with MED1 or MED26 occupancy did not colocalize (**Figure 3A and Fig. S5A**). Notably, MED26 colocalizes with GATA1 and GATA2 better than MED1, indicating a potential functional association between MED26 and GATA factors that could play a pivotal role in the regulation of erythroid transcriptome (**Figure 3A**). The presence of MED1-or MED26-enriched puncta were observed in primary human erythroblasts as well as other cell types (**Fig. S5B-5C**). We then calculated the signal ratio of MED26 to MED1 at the transcription start sites (TSSs) of their occupancy loci. While a large fraction of MED1- and/or MED26-occupied genes have a relatively constant MED26/MED1 signal ratio, a considerable number of genes have differential MED26/MED1 ratios, indicating that the chromatin occupancy of MED26 and MED1 does not always correlate. We defined the maximum 10% ratio as an indicator of MED26-enriched chromatin sites and the minimum 10% ratio as an indicator of MED26-poor chromatin sites (**Figures 3B-3C**). To elucidate how the enrichment of MED26 impacts transcription, we conducted a CUT&Tag assay of Pol II and precision nuclear run-on sequencing (PRO-seq) in human CD34^+^ erythroid culture cells on Day 4. We found that Pol II has more signals spanning through the gene body at MED26-poor chromatin sites, whereas Pol II often has distinct enrichment at the TSS region, transcribing a higher percentage of short RNAs at MED26-enriched chromatin sites and indicating Pol II pausing (**Figures 3D-3E**). To better characterize this observation, we calculated the pausing index (PI), which is defined by the ratio of the read count at the TSS region to the read count at the gene body region after normalizing both read counts by the length of the genomic region (**Fig. S5D**)^25^. Our analysis revealed that the MED26-enriched loci had a significantly higher PI (p < 2.2 × 10^-16^), longer gene length (p = 4.092 × 10^-10^), and similar exon numbers (p = 0.09774) than the MED26-poor loci (**Figures 3F and Fig. S5E**). Based on the overall quantification, we found that Pol II in MED26-enriched chromatin sites transcribes genes with a higher PI, lower elongation efficiency and a higher percentage of short RNAs (**Fig. 3G**).

**Figure 3.**
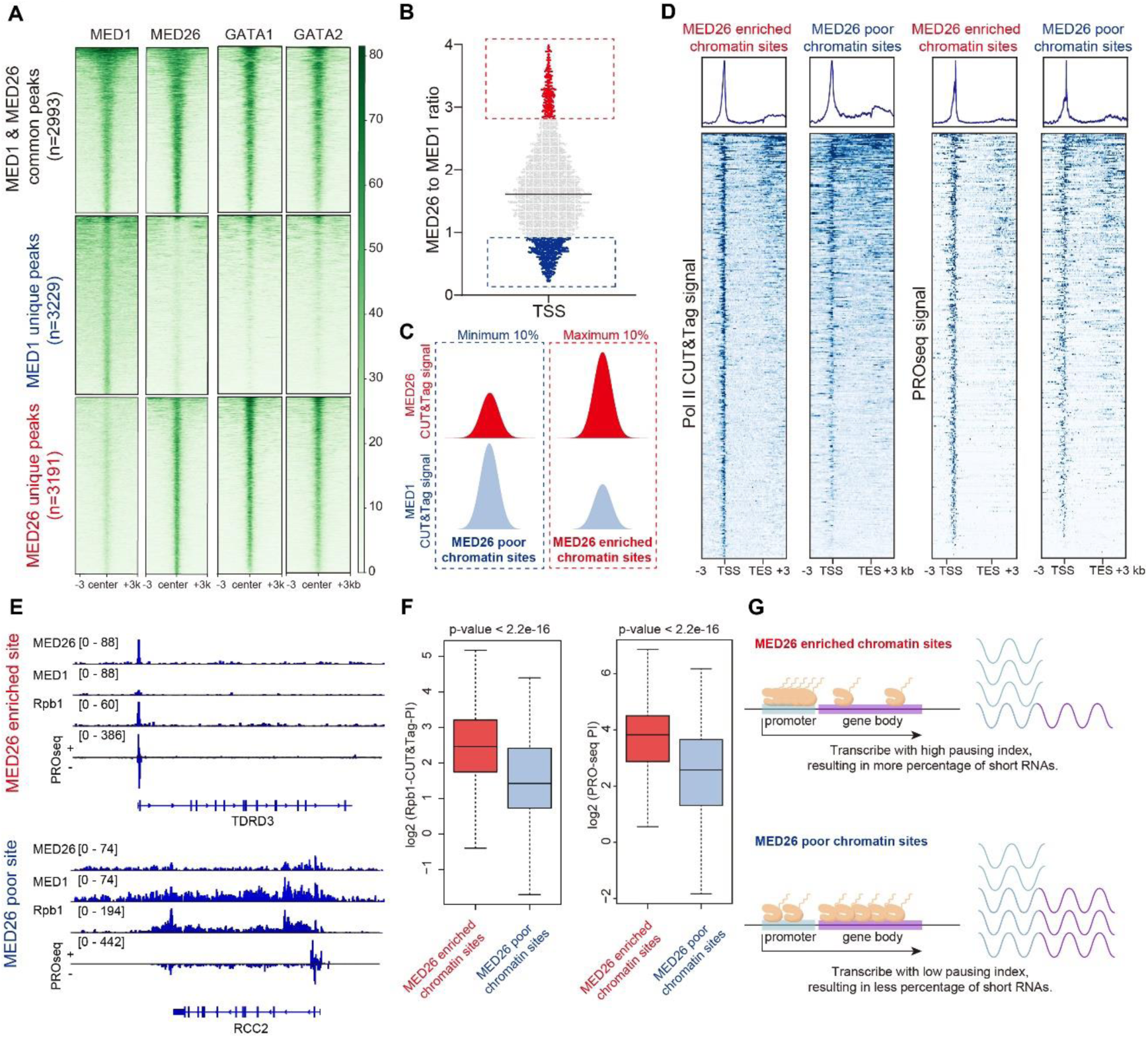
RNA polymerase Ⅱ exhibits transcription pausing at MED26-enriched loci. **(A)** Heatmaps from CUT&Tag assays showing the number and distribution of MED1, MED26, GATA1 and GATA2 common peaks; MED1 unique peaks, and MED26 unique peaks on Day 4 in the primary human CD34^+^-derived erythroid culture. **(B)** The ratio of MED26 to MED1 signals on all the transcription start sites (TSSs) with MED1 and/or MED26 occupancy in CUT&Tag assays of primary human erythroblasts on Day 4. Each dot represents one gene. **(C)** Schematic defining the MED26-enriched or MED26-poor Mediator loci. **(D)** Heatmaps showing RNA Pol Ⅱ and PRO-seq signal of MED26-enriched or MED26-poor genes from −3 kb of the TSS to the +3 kb of transcription end site (TES). **(E)** IGV visualization showing examples of MED26-enriched (top) and MED26-poor (bottom) loci. RPB1 is the largest component of RNA Pol Ⅱ. **(F)** Boxplots comparing the pausing indices of MED26-enriched or MED26-poor loci. The pausing indices were calculated from RPB1 CUT&Tag or PRO-seq. The p values were calculated using the two-sided Wilcoxon rank-sum test. **(G)** Model representing the different transcription behaviors of MED26-enriched or MED26-poor condensates.

### MED26 recruits pausing-related factors to mediate transcription pausing

To examine how MED26 might mediate transcription pausing, we performed immunoprecipitation coupled with mass spectrometry (IP-MS) to identify interacting proteins of MED1 and MED26 and then analyzed the relative protein interaction using normalized spectral abundance factors (NSAFs)^26^. While MED26 and MED1 similarly coimmunoprecipitated with the majority of other Mediator subunits (the head, middle, and tail modules) and the elongation complex, MED26 interacted much less with the CDK8 kinase module than MED1, consistent with a previous study^27^. Remarkably, MED26 exhibited a substantially higher interaction with transcription pausing related factors (NELF, DSIF, and PAF complexes) compared to MED1, which supports our notion that MED26 associates with pausing (**Figures 4A-4B and Table S1**). Knocking out MED26 significantly impairs the recruitment of PAF1 to MED26 occupancy sites, indicating the presence of MED26 facilitates the recruitment of PAF1 (**Figure 4C**). Promoter-proximal pausing of Pol II after initiation represents a poised stage which requires additional activation for productive transcription elongation^14,28^. Our findings, demonstrating the comparable interactions of MED26 with elongation factors and its markedly stronger interactions with pausing-related factors compared to MED1, may explain their differences in regulating Pol II behavior.

**Figure 4.**
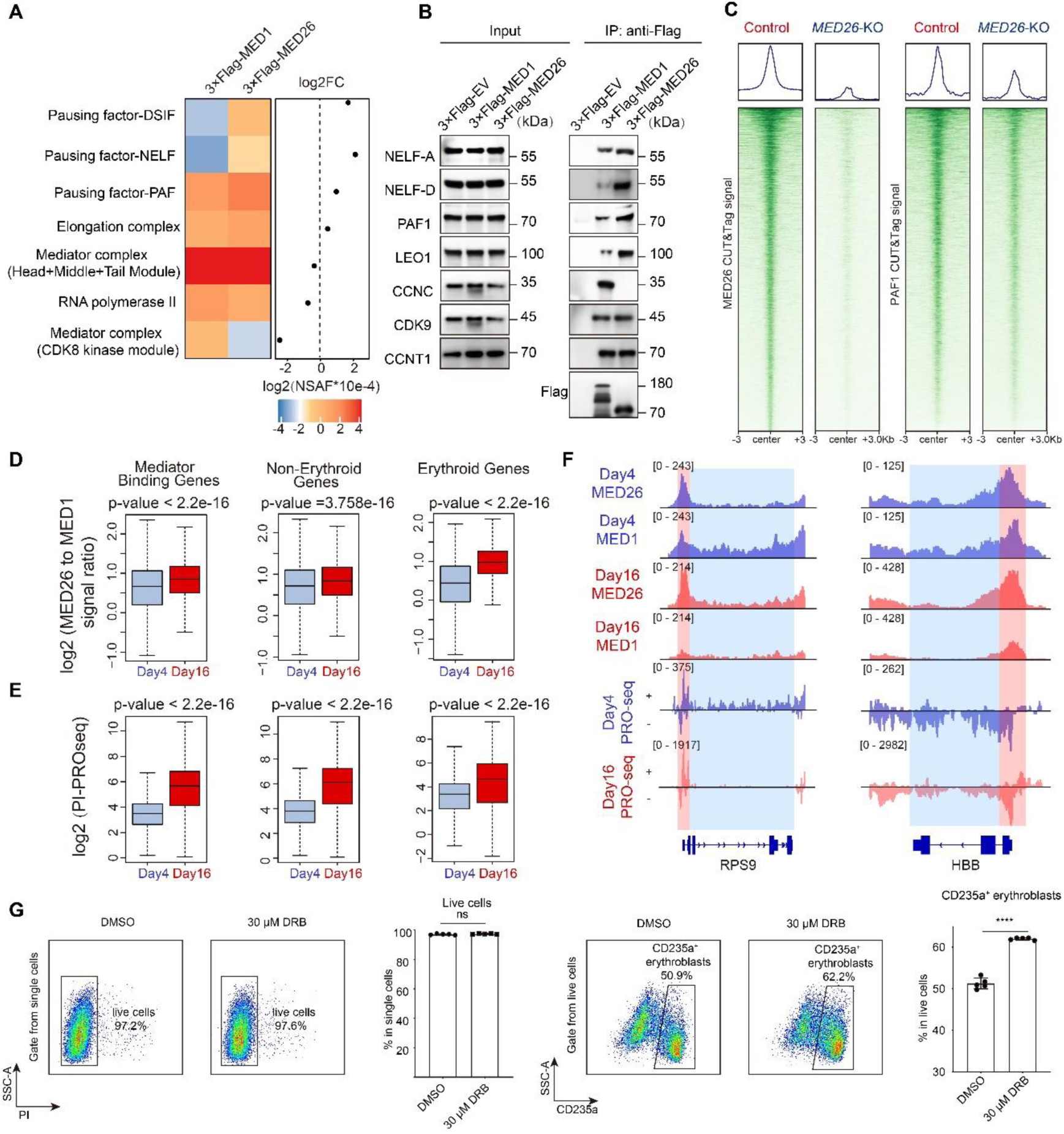
Transcription pausing is essential for erythropoiesis. **(A)** Heatmap showing the relative interaction strengths of the indicated proteins baited with MED1 or MED26 in HEK293 cells. The relative interaction strength was calculated according to the NSAFs from mass spectrometry data (*see Methods*). **(B)** Coimmunoprecipitation of HEK293 cells transfected with 3×Flag-MED1, 3×Flag-MED26, or 3×Flag-EV (control plasmid). Following immunopurification by Flag antibody, western blotting was performed on the complexes using antibodies against components of the pausing complexes (NELF-A, NELF-D, PAF1, and LEO1), the Mediator CDK8 kinase module (CCNC1), and the elongation complex (CDK9 and CCNT1). **(C)** Heatmaps showing MED26 and PAF1 signal distribution of −3∼+3 kb regions around peak centers in wild type or *MED26* KO K562 cells. **(D-E)** Boxplots comparing the MED26 to MED1 signal ratio and the PI on Day 4 and Day 16 of the human CD34^+^-derived erythroid culture. Day 4 *ex vivo* cultured cells primarily consisted of HSPCs and some erythroid progentiors, while day 16 cells predominantly included late erythroblasts. (The p values are calculated by paired Wilcox test. Nonerythroid genes and erythroid genes were defined based on published data^50^). **(F)** IGV visualization of the CUT&Tag signals of PRO-seq, MED1, and MED26 at the *RPS9* (nonerythroid gene) and *HBB* (erythroid gene) loci. **(G)** FACS analysis detecting erythroid differentiation upon treatment with 30 µM DRB (5,6-dichloro-1-beta-D-ribofuranosylbenzimidazole) in Day 6 primary human erythroblasts, performed on Day 10. (Left) Cell death rate after DRB treatment, assessed by PI staining. (Right) Accelerated erythroid differentiation detected by the percentage of CD235a-positive cells. Data are represented as the mean ± SD (n=5). The p values were calculated using an unpaired two-tailed Student’s *t* test, and significant differences are marked by asterisks: ****p<0.0001, n.s., not significant.

### MED26-mediated transcription pausing serves as a critical mode of gene regulation in terminal erythropoiesis

To address how the enrichment of MED26 impacts transcription during erythropoiesis, we used CUT&Tag assays to analyze the ratio of MED26 to MED1 chromatin occupancy on Day 4 and Day 16 of primary human CD34^+^-derived erythroblasts. Our results revealed that the ratio of MED26 to MED1 increased at both erythroid and nonerythroid genes during differentiation, and the ratio at erythroid genes on Day 16 had a more marked increase relative to that at nonerythroid genes (**Figure 4D**). We then performed PRO-seq on Day 4 or Day 16 *ex vivo*-cultured human erythroblasts. Our results revealed that the PI increased at both nonerythroid genes (p < 2.2 × 10^-16^) and erythroid genes (p < 2.2 × 10^-16^), along with an increase in the MED26 to MED1 ratio during erythroid development (**Figure 4E**). Non-erythroid gene *RPS9* showed decreased occupancy of MED1 and MED26 from Day 4 to Day 16, with a more pronounced decrease of MED1 signal; erythroid gene *HBB* showed increased occupancy of MED1 and MED26 from Day 4 to Day 16, with a more profound increase of MED26 signal (**Figure 4F**). Meanwhile, a higher percentage of nascent short RNAs were transcribed and accumulated at both non-erythroid and erythroid genes, resulting from transcription with a higher pausing index and lower elongation efficiency in terminal erythropoiesis (**Figure 4F**). To test whether the increase of transcription pausing accelerates erythropoiesis, we added 5,6-dichloro-1-β-D-ribofuranosylbenzimidazole (DRB), which represses transcription elongation by inhibiting CDK9^29^, to Day 6 *ex vivo*-cultured human erythroblasts. The addition of DRB increased the number of CD71^+^CD235a^+^ erythroblasts compared to the mock group (**Figure 4G**). Collectively, our findings reveal that MED26 enrichment is associated with increased transcription pausing during terminal erythroid stages, which can accelerate erythroid development.

### The condensate-forming capacity of MED26 is essential for erythropoiesis

Based on previous structural studies^30,31^, MED26 contains three domains— a TFIIS domain (amino acids (a.a.) 1-87), an unstructured domain (a.a. 88-480), and a Mediator complex interacting domain (a.a. 480-600). To determine which domain is important for the erythropoiesis promotion function of MED26, we overexpressed these three segments of MED26 in human CD34^+^ HSPCs, which showed that a.a. 88-480 alone can sufficiently accelerate erythroid differentiation as full-length MED26 (**Figure 5A**). As unstructured low-complexity domains tend to undergo phase separation, we used the OptoDroplet assay, in which proteins with phase separation capacity will aggregate when triggered by blue light, to examine MED26 truncations (**Fig. S6A**). The results showed that the truncations containing a.a. 88-480, the putative intrinsically disordered region (IDR), exhibited aggregate-forming capacity (**Fig. S6B-6C and Movie S1-4**). Interestingly, a.a. 1-480 formed more conspicuous aggregates than all other truncations, which suggests that a.a. 1-87 can promote the phase separation capacity of the MED26 IDR (**Fig. S6B-6C and Movie S1-4**). *In vitro* assays further validated that a.a. 88-480 of MED26, but not a.a. 1-87 or 480-600, had phase separation capacity (**Figure 5B**). Collectively, these results indicate that a.a. 88-480, the phase-separation domain of MED26, is the key segment promoting erythroid differentiation.

**Figure 5.**
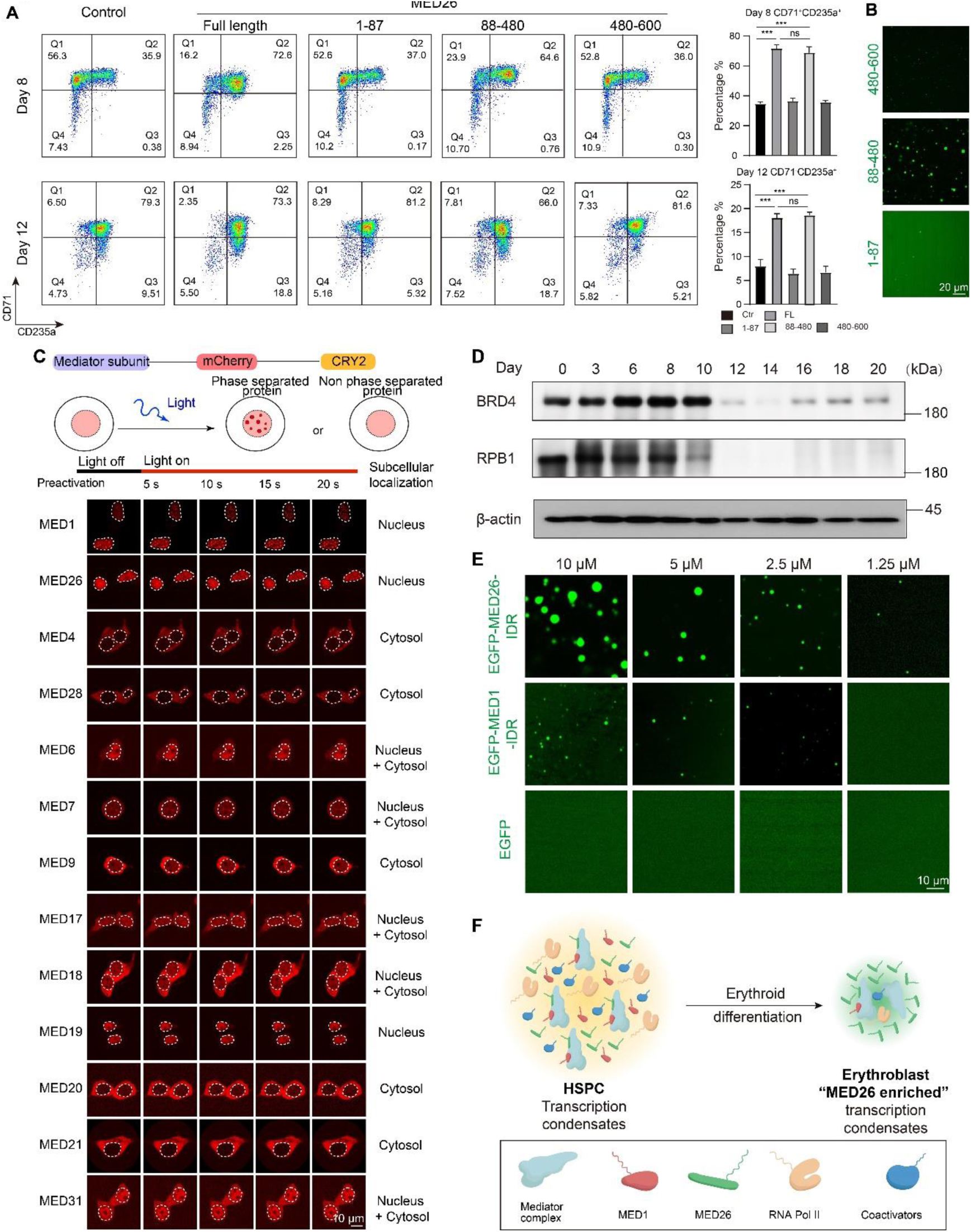
MED26 exhibits a strong condensate-forming capacity and its condensate-forming domain is sufficient to accelerate erythroid differentiation. **(A)** FACS analysis to detect erythroid markers (CD71, CD235a) upon full-length or truncated MED26 overexpression in *ex vivo* human CD34^+^ erythroid culture. Data are represented as the mean ± SD (n=3). The p values were calculated using an unpaired two-tailed Student’s *t* test, and significant differences are marked by asterisks: ***p<0.001, n.s., not significant. **(B)** *In vitro* phase separation assay of the indicated MED26 truncations fused with EGFP in a low salt buffer. **(C)** (Top) Schematic diagram of the OptoDroplet assay. The candidate protein was fused with mCherry and CRY2. Upon blue light stimulation, only disordered proteins can form aggregates. (Bottom) Time-lapse images of various Mediator subunits analyzed by the OptoDroplet assay. The subunits are indicated on the left, and subcellular localization is indicated on the right. **(D)** Western blot showing the relative abundance of BRD4 and RPB1 over time (Days 0-20) in human CD34^+^ erythroid culture with β-actin as a loading control. **(E)** Droplet formation assays of gradient-diluted MED26 and MED1 IDR proteins in low-salt buffer without PEG. **(F)** Model representing transcription condensate component switching during erythroid differentiation.

### MED26 exhibits more pronounced capability of biomolecular condensate formation than MED1

Biomolecular condensates have emerged as a prevalent mechanism for transcription control^32^. Intrinsically disordered regions (IDRs) and/or low-complexity protein sequences are necessary for biomolecular condensate-mediated phase separation and are predicted to widely exist in Mediator subunits^33^. MED1 has been reported to mediate transcription regulation via phase separation and serves as a surrogate to study phase separation of transcription machinery^34,35^, but it remains unclear whether the other Mediator subunits also form liquid-like condensates. Therefore, in addition to MED26, we also tested whether other Mediator subunits have condensate-forming capability by conducting the OptoDroplet assay (**Figure 5C**)^36^. This assay revealed that MED1, MED4, MED26, and MED28 had droplet-forming capability (**Figure 5D and Movie S5-17**), consistent with the prediction by natural disordered region (PONDR) analysis (**Fig. S7A**) and structural evidence showing that large parts of these proteins are unstructured^30^. During erythropoiesis, we not only found that the level of most Mediator subunits declined but that of representative transcription condensate-forming proteins, such as BRD4 and RPB1, also diminished substantially (**Figures 1A-1C and 5D**). Taken together, these findings revealed a distinct stoichiometric composition of individual Mediator subunits in transcription condensates during terminal erythroid differentiation.

We examined whether MED26 exhibits droplet properties *in vivo* by the fluorescence recovery after photobleaching (FRAP) assay, which showed that enhanced green fluorescence protein fused with MED26 (EGFP-MED26) can form condensates in K562 erythroleukemia cells and that the signal recovers promptly after photobleaching (**Fig. S7B and Movie S18**). Immunofluorescence imaging showed that Mediator condensates were diffused upon treatment with 1,6-hexanediol (1,6-HD), which can disrupt liquid-like condensates (**Fig. S7C**). To better characterize the intrinsic condensate-forming capacity of MED26, we expressed and purified putative EGFP-MED26-IDR (**Fig. S7D**) ^30^. We observed that EGFP-MED26-IDR can form phase-separated droplets *in vitro*, with or without crowding reagents (10% PEG8000) (**Fig. S7E-7G**). Using time-lapse imaging, we were able to capture fusion events of the MED26 droplets, indicating their fluidic property (**Fig. S7E and Movie S19-20**). We then performed a droplet formation assay with varying concentrations of EGFP-MED1-IDR, EGFP-MED26-IDR, and EGFP, which showed that MED26-IDR has a lower saturation concentration of phase separation than MED1-IDR (**Figure 5E**). Moreover, we observed that MED26-IDR droplets are sensitive to 1,6-HD and high salt treatment, indicating that hydrophobic and electrostatic interactions contribute to droplet formation (**Fig. S7F-7G**). Collectively, our results demonstrate that MED26, a subunit in the core Mediator complex, can undergo phase separation *in vitro* and *in vivo* and the transcription condensates shift towards a MED26-enriched form during terminal erythroid differentiation (**Figure 5F**).

### The erythropoiesis-promotion function of MED26 is dependent on its capacity of recruiting pausing-related factors

To determine whether the recruitment of pausing-related factors depends on the phase separation capacity of MED26, we purified mCherry fused with the disordered region of a pausing factor, PAF1. The *in vitro* droplet formation assay not only showed that a.a. 1-480 of MED26 can recruit PAF1, but also suggested that the presence of pausing-related factors reciprocally promotes condensates formation of MED26 (**Figure 6A**). To investigate whether the IDR of MED26 contributes to the recruitment of pausing-related factors, we constructed EGFP fused with various truncations of MED26 and tested the recruitment of PAF1-mCherry. The results demonstrate that MED26’s ability to recruit PAF1 is contingent on its IDR, which cannot be substituted by the FUS IDR domain, a known sequence that can drive liquid-like condensate formation (**Figure 6B**). These observations imply that MED26’s IDR domain plays a distinctive role in recruiting PAF1. The PAF1 complex could regulate promoter proximal pausing and/or release depending on the cellular context, but PAF1C was initially regarded as a pausing factor, akin to the DSIF complex, that counteracts transcription elongation in erythropoiesis^37–40^. Knocking down PAF1 or SPT5 (a known pausing factor, subunit of DSIF complex) in the human CD34^+^ erythroid system similarly inhibits erythropoiesis, indicating a crucial role of pausing in erythropoiesis (**Figure S8A-8B**). To further investigate the functional association between MED26 and pausing-related factors in erythropoiesis, we conducted simultaneous PAF1 knockdown and over-expression of MED26 in the primary human CD34^+^ erythroid culture system. The results revealed that the erythropoiesis promotion by MED26 was abolished by PAF1 knockdown (**Figures 6C-6E**). Taken together, the findings indicate that both the IDR and TFIIS domains of MED26 are necessary for the recruitment of pausing-related factors to the MED26-enriched condensates, and MED26 accelerates erythropoiesis via its association with these factors.

**Figure 6.**
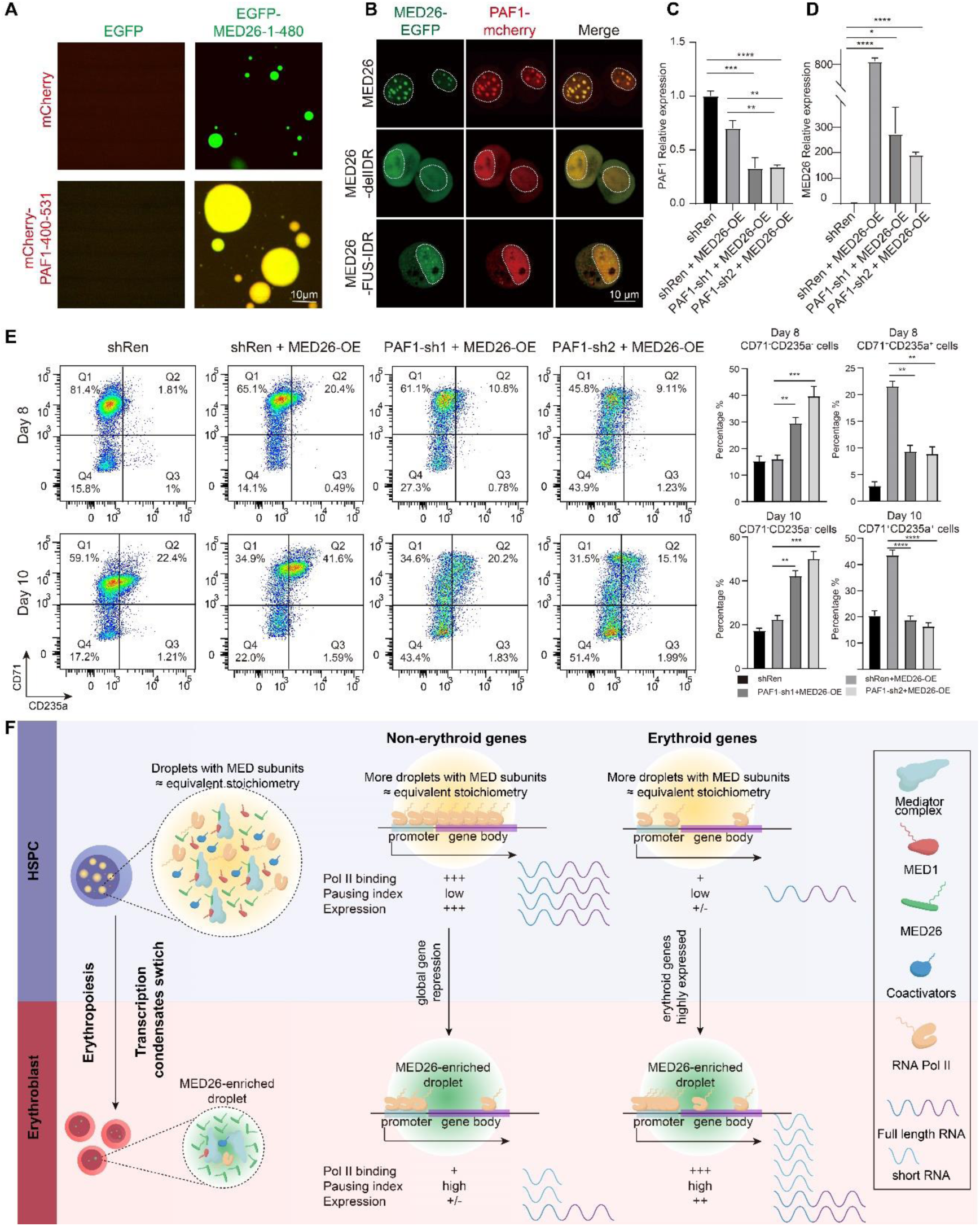
MED26 accelerates erythropoiesis by recruiting pausing factors through its phase separation domain. **(A)** *In vitro* phase separation assay of EGFP-fused MED26 1-480 and mCherry-PAF1-400-531 with a buffer containing 5% PEG-8000. **(B)** Representative images of *MED26*-KO K562 cells transfected with MED26-truncations-EGFP and PAF1-mCherry. The white dotted circle indicates the nuclear area. **(C-D)** qPCR validation of the relative expression of PAF1and MED26 in indicated CD34^+^ cells. qPCR data are represented as the mean ± SD (n=3). **(E)** FACS analysis to detect two erythroid markers (CD71 and CD235a) upon simultaneous PAF1 knockdown and MED26 over-expression. Data are represented as the mean ± SD (n=3). **(F)** Model representing the transcription features of erythroid and nonerythroid genes during different differentiation stages. The p values were calculated using an unpaired two-tailed Student’s *t* test, and significant differences are marked by asterisks: *p<0.05, **p<0.01, ***p<0.001, ****p<0.0001.

Transcription pausing is an important mode of transcription regulation that is often associated with rapid and synchronous gene induction in development^13^. To understand how MED26-mediated transcription pausing regulates terminal erythroid differentiation, we analyzed overall gene expression and Pol II occupancy during erythroid differentiation. We found that the expression of nonerythroid genes was downregulated, while that of erythroid genes was upregulated (**Supplemental Figure 8C**). Furthermore, we found that Pol II occupancy decreased at nonerythroid gene loci, while that at erythroid loci increased during erythroid development (**Supplemental Figure 8D**). Taken together, the data suggest that there is an overall increase in transcription pausing resulting from the shift of transcription condensates towards a “MED26 enriched form”, leading to decreased elongation efficiency and global transcription repression during terminal erythropoiesis. Under the condition of increased pausing, nonerythroid genes exhibit low Pol II occupancy, which results in decreased gene expression; erythroid genes have higher Pol II occupancy, despite exhibiting an accumulation pattern at the TSS, still resulting in the production of abundant full-length transcripts (**Figure 6F**).

## Discussion

Global gene repression is a unique and pivotal feature of terminal erythropoiesis, the mechanism of which remains elusive^4^. MED1, the representative subunit of the Mediator complex, was shown to be critical for erythropoiesis *in vivo*^19,20^, but it remains unclear whether other subunits exert specific functions during this process. In this study, we uncovered that the composition of Mediator-containing transcription condensates are highly dynamic during erythropoiesis, with most subunits diminishing while MED26 remained relatively abundant throughout the process. By comparing the chromatin occupancy sites of MED26 and MED1, we revealed that MED26-enriched chromatin sites are associated with transcription pausing. We uncovered that a major unstructured segment (IDR) of MED26 is necessary for its function in promoting erythropoiesis. MED26 has a much stronger preference to interact with pausing-related factors compared to MED1. Moreover, the IDR of MED26, which exhibits pronounced condensate forming capability, is essential for recruiting pausing-related factors to the MED26-enriched condensates. Altogether, our findings demonstrate that the relative enrichment of MED26 in terminal erythropoiesis leads to a shift of the transcription condensates towards a “MED26-enriched form”. This shift, in turn, facilitates the recruitment of pausing-related factors, thereby achieving global gene repression.

Biomolecular condensates-mediated dynamic cellular compartmentalization has emerged as a common mechanism in various aspects of transcription regulation^32^. Recently, a number of transcription regulatory proteins, including MED1, BRD4, and Pol II CTD^34,35,41^, have been reported to form biomolecular condensates or membraneless compartments mediated by phase separation, which has become an emerging model to explain diverse cellular events, including transcription regulation^42–44^. The presence of promoter condensates and gene-body condensates has been proposed at different transcription steps^45,46^. Nonetheless, it remains unclear whether the dynamic composition of these condensates could contribute to driving the progression of developmental processes. Our study not only found that MED26 exhibits more conspicuous condensate-forming capacity than most other subunits, but also identified MED26-enriched condensates as a dominant form of transcription condensates in terminal erythropoiesis, prominently facilitating Pol II pausing, which is pivotal for global transcription repression of terminal erythropoiesis.

Transcription pausing can serve as an important mechanism for precise transcription regulation^14^. Regarding how erythroid genes remain transcriptionally active in terminal erythropoiesis, we propose two hypotheses. First, the master erythroid transcription factors GATA1 and EKLF have been shown to recruit P-TEFb, which enhances transcription elongation^8^. Moreover, erythroid genes are enriched with elongation factors. In zebrafish erythropoiesis, TIF1γ-dependent recruitment of positive elongation factors to erythroid genes counteracts Pol II pausing^37^. Second, critical cytokines such as erythropoietin (EPO) can stimulate pause release^47^. Therefore, although MED26 has a preference for recruiting pausing-related factors, TFs or cytokines could, in turn, relieve transcription pausing. Therefore, MED26 may serve as a gatekeeper for pausing Pol II, which may remain paused until further signaling for productive elongation to occur.

All in all, our study utilizes erythropoiesis as a unique paradigm to provide mechanistic insights into context-dependent transcription regulation during development. We have uncovered that the composition of Mediator-containing transcription condensates can be highly dynamic, and different forms of transcription condensates contribute to the versatile transcriptional and functional outputs. Our study identifies a stoichiometric switch of Mediator during erythroid development, and MED26-enriched condensates serve as an important regulator for transcription pausing and could potentially be a therapeutic target for diseases, such as erythroid dysplasia.

## Methods

Detailed methodology is provided in the Supplemental information file.

### *Ex vivo* CD34^+^ cell erythroid differentiation system

CD34^+^ cells were purified from human cord blood (Cord Blood Bank of Beijing) using the MACS MicroBead kit (Miltenyi Biotec, Bergisch Gladbach, Germany). The two-phase erythroid differentiation protocol was modified from the previous study^23^.

### Mice

All mice were constructured by CRISPR-Cas9 gene-editing technology and bred under specific-pathogen-free (SPF) conditions. All mouse procedures were approved by the Animal Care and Use Committees of Peking University. The detailed information about mice are provided in the Supplemental information file.

### High-throughput sequencing library construction and analysis

The CUT&Tag assay was performed as previously described^48^ and followed the manufacturer’s instructions for the kit (Vazyme, Nanjing, China). PRO-seq libraries were prepared as described previously^49^. The detailed information about high-throughput sequencing library construction and analysis are provided in the Supplemental information file.

### Data sharing statement

All sequencing data are available in the NCBI Gene Expression Omnibus (GEO) database (GSE212374). The code is available upon request.

## Acknowledgments

We thank the Flow Cytometry Core and the Imaging Core of the National Center for Protein Sciences at Peking University (PKU). We thank the High-Performance Computing Platform of the Center for Life Sciences (Peking University) for supporting data analysis. We thank Drs. Zhi Qi, Xiong Ji, Jiazhi Hu and Chengqi Yi (PKU) for their suggestions on the project. We are grateful to Dr. Hongxia Lv, Dr. Liying Du, Dr. Chunyan Shan, Dr. Siying Qin, Dr. Dong Liu, Dr. Qi Zhang, Dr. Guilan Li, Dr. Jia Luo, Huan Yang, Xuefang Zhang, Yinghua Guo, and Jun Cheng (PKU) for technical support. We thank all the other members of the laboratory for their critical reading of this manuscript. This project was supported by grants from the National Key R&D Program of China (2022YFA1103300), the National Natural Science Foundation of China (82370124), and the Peking-Tsinghua Center for Life Sciences and School of Life Sciences, Peking University, to H.Y.L. and by grants from Department of Science and Technology of Zhejiang Province (Key Program Grant, 2021YFA1100100), the National Natural Science Foundation of China (81973993), the Zhejiang Provincial Natural Science Foundation of China (LR20C070001), and the Hangzhou Science and Technology Major Project (2018HZKJSA10095) to X.G.

## Authorship contributions

S.Z., X.G. and H.Y.L. conceptualized this project and designed the experiments. S.Z., M.L., X.Y.Z., S.X., Q.Y., N.L., Y.L., D.L., Z.L.Z., Z.F., X.J., Y.D., Q.H., Y.Y.Z. and W.H.C. performed the experiments. S.Z., X.T.Z., X.Y.Y. and F.W. conducted the bioinformatic analyses. S.Z., X.G. and H.Y.L. wrote the manuscript. All authors discussed the results and commented on the manuscript.

